# Characterizing The Multimodal Sympathetic Nervous System Startle Response

**DOI:** 10.64898/2026.03.13.711638

**Authors:** Ramanamurthy V. Mylavarapu, Eric R. Albuquerque, Gary J. Farkas, David W. McMillan, Patrick D. Ganzer

**Affiliations:** Department of Biomedical Engineering, University of Miami, Coral Gables, FL; The Miami Project to Cure Paralysis, University of Miami Miller School of Medicine, Miami, FL; Medical Scientist Training Program, University of Miami Miller School of Medicine, Miami, FL; Department of Physical Medicine and Rehabilitation, University of Miami Miller School of Medicine, Miami, FL; Department of Neurological Surgery, University of Miami Miller School of Medicine, Miami, FL

## Abstract

The human startle reflex has primarily been characterized by its repeatable, coordinated, and temporally patterned somatomotor responses. In this study, we assessed whether startle-evoked sympathetic nervous system (SNS) responses might also constitute a repeatable, coordinated, and temporally structured output. Using a noninvasive tactile startle stimulus, we simultaneously recorded startle-evoked electrodermal activity, photoplethysmography-derived blood volume indices, heart rate, blood pressure, stroke volume, and cardiac output in healthy participants. Our results demonstrate that startle elicits a reproducible and patterned constellation of SNS responses – a multimodal ‘SNS startle signature’ – with conserved temporal relationships across effector systems. The SNS startle signature was composed of robust bilateral peripheral responses, including palmar sweating, cutaneous vasoconstriction, and biphasic cutaneous venous-capillary blood volume changes, in addition to more mild central hemodynamic changes. In contrast to previous startle reflex studies, there was no influence of biological sex or cardiac-cycle gating on responses. Lastly, the SNS startle signature exhibited features of a potentially attractive diagnostic, associated with robust responder discrimination and high response reliability across repeated trials. Overall, these findings fill a critical knowledge gap and also suggest the potential utility of this multimodal signature for assessing autonomic dysfunction.

## 1. INTRODUCTION

The sympathetic nervous system (SNS) plays a central role in rapidly coordinating responses to novel or threatening challenges^1^. SNS activation can originate from top-down cortical control and reflexive brainstem-spinal pathways, and includes the startle reflex^2,3^. The startle reflex rapidly coordinates responses from multiple systems in response to a threatening stimulus. These include: (i) skeletal muscle^4–8^, (ii) sweat gland^9–12^, (iii) vasomotor^13,14^, and (iv) cardiovascular responses^12,15–17^. Historically, the human startle reflex has primarily been defined and quantified through its somatomotor manifestations, specifically eyeblink, craniofacial, and postural muscle responses^4–8^. However, despite the clear involvement of the SNS during startle, SNS responses have comparatively received far less attention, specifically regarding their timing, coordination, and possible patterned outputs.

Startle-triggered somatomotor responses require measurement from the skeletal muscle system, and are characterized by highly patterned latencies, amplitudes, and coordination across muscle groups^18–21^. In contrast, SNS responses evoked by startle affect multiple systems, have typically been studied in isolation, and are commonly treated as unstructured indices of arousal^22–25^. Furthermore, even fewer studies have assessed multiple sympathetic effector systems simultaneously. Consequently, it remains unclear whether startle-evoked SNS responses represent a diffuse unstructured arousal reaction, or instead constitute a coordinated temporally patterned output analogous to the well-established somatomotor startle response.

Lastly, similar to other reflexes, a multimodal SNS startle response may also have utility as a diagnostic tool for assessing autonomic nervous system (ANS) dysfunction. ANS dysfunction is widely implicated in neurologic, cardiovascular, and metabolic diseases. The clinical measures for characterizing ANS dysfunction can either be difficult to apply or involve tools that only assess a single SNS process alone^9,26–29^. For example, the Composite Autonomic Severity Score (CASS) and Quantitative Sudomotor Axon Reflex Test (QSART) are both widely used clinical measures for assessing ANS dysfunction^30^. However, these tools rely on specialized or invasive procedures and only provide an incomplete assessment of distributed SNS function. In addition, sweat response changes (e.g., measured by skin conductance) in response to a short noninvasive electrical stimulus has also been used for the detection of peripheral nerve dysfunction following both spinal cord injury (SCI)^9,26,28^ and diabetic peripheral neuropathy^31,32^. Regardless, these approaches only focus on a single SNS process. In addition, several other clinical studies have leveraged the startle reflex to assess function in patients with disease (e.g., Parkinson’s disease^33,34^, stroke^35,36^, post-traumatic stress disorder (PTSD)^37,38^, and schizophrenia^39,40^). However, these approaches overwhelmingly focus on somatomotor responses alone and not SNS processes. Overall, a distributed and structured SNS startle response may have utility for evaluating ANS function in clinical populations. Importantly, establishing the temporal structure and coordination of startle-evoked SNS responses in healthy subjects is necessary prior to applications in clinical populations with autonomic disease and dysfunction.

In this study, we assessed two main hypotheses: 1) startle-evoked SNS responses generate a highly structured and patterned multimodal SNS startle signature, and 2) the SNS startle signature can potentially be used as an ANS diagnostic measure, associated with robust responder discrimination and high response probability. In healthy subjects, the startle reflex was triggered using a noninvasive electrocutaneous stimulus, and electrodermal (EDA), photoplethysmography (PPG) derived blood volume indices, stroke volume (SV), cardiac output (CO), heart rate (HR), and mean arterial pressure (MAP) signals were recorded noninvasively. In addition, similar to other ANS diagnostic measures, we also characterized responder profiles, response habituation, and the predictive performance for startle-triggered SNS responses. Overall, our results characterize the dynamics and diagnostic potential of the SNS startle signature, demonstrating its repeatable, highly coordinated, and patterned output.

## 2. METHODS

### 2.1 Study Participants

This study was approved by the Institutional Review Board (IRB) of the University of Miami (IRB# 20190769). Informed consent was obtained from all study participants before any study procedures were performed. Fifteen healthy adults participated in this study (Table 1). All participants had no pre-existing cardiovascular, neurological, or rheumatological diseases.

**Table 1.** Summary of Participant Data and Baseline Measures.

|  | Total (n=15) | Male (n=7) | Female (n=8) | p-val* |
| --- | --- | --- | --- | --- |
| Age (y) | 26.67 ± 4.84 | 26.71 ± 4.27 | 26.63 ± 5.40 | 0.98 |
| Ambient Room Temp (°C) | 21.45 ± 1.02 | 21.54 ± 0.89 | 21.35 ± 1.15 | 0.72 |
| Participant Limb Temp (°C) | 31.79 ± 2.21 | 32.26 ± 2.03 | 31.31 ± 2.38 | 0.42 |
| Systolic BP (mmHg) | 109.56 ± 12.60 | 112.86 ± 10.78 | 106.25 ± 14.41 | 0.33 |
| Diastolic BP (mmHg) | 75.92 ± 7.48 | 73.71 ± 4.68 | 78.13 ± 10.27 | 0.30 |
| HR (bpm) | 68.32 ± 9.91 | 63.00 ± 11.03 | 73.63 ± 7.35 | 0.06 |

### 2.2 Experimental Paradigm Overview

All experimental sessions were conducted in the morning, between 8:00 and 10:30 AM. Participants were instructed to fast and refrain from consuming caffeinated beverages 12 hours before the session. The morning of the recording, participants sat in the experiment room at rest for at least 10 minutes to acclimate to the room temperature (19 – 23 °C). The Perceived Stress Scale (PSS-10)^41^, Generalized Anxiety Disorder Questionnaire (GAD-7)^42,43^, Beck Depression Inventory (BDI)^44^, and Adverse Childhood Experience Questionnaire (ACEs)^45^ were completed by all participants. We also recorded the participants’ self-reported hours of sleep, exercise performance, and abstinence from food and caffeinated beverages prior to the session. Otherwise, participants were instructed to not alter their normal morning routine or normal medication regimen, if applicable. Of note, one of the participants was on medications when enrolled in the study. Further analysis revealed that their recorded signals were similar to the overall participant cohort, and their data were included in the final analyses. Once the acclimation period and questionnaires were completed, noninvasive sensors measuring skin conductance, peripheral vasomotor activity, and cardiovascular parameters were placed on the participant (**Figure 1A**). Skin temperature from the right palm, blood pressure (BP), and HR were measured before the start of the session. The recording session consisted of 54 trials in total: 3 initial trials of no nerve stimulation, followed by 48 trials of median nerve stimulation, and finally 3 additional trials of no nerve stimulation. Each stimulation trial consisted of a soft low-volume auditory cue approximately 30 seconds before nerve stimulation, signaling participants to passively look forward during stimulus delivery. Signals were recorded for 20 seconds before and 50 seconds after each stimulation. The first and last 10 seconds of every trial were used as a recording buffer and were removed prior to all data analyses (this prevented introducing equipment or participant artifacts at the beginning and end of trials). This trial structure was informed overall by preliminary data and previous studies suggesting that a 1- to 2-minute washout period between stimulation trials is sufficient to avoid confounding effects of habituation or trial bleed-through^28,46^. A second skin temperature measurement was obtained at the end of the session, as peripheral skin temperatures have been shown to affect sudomotor and vasomotor signals^47,48^. Please see **Supplemental Figure S3** for more details on temperature readings.

**Figure 1.**
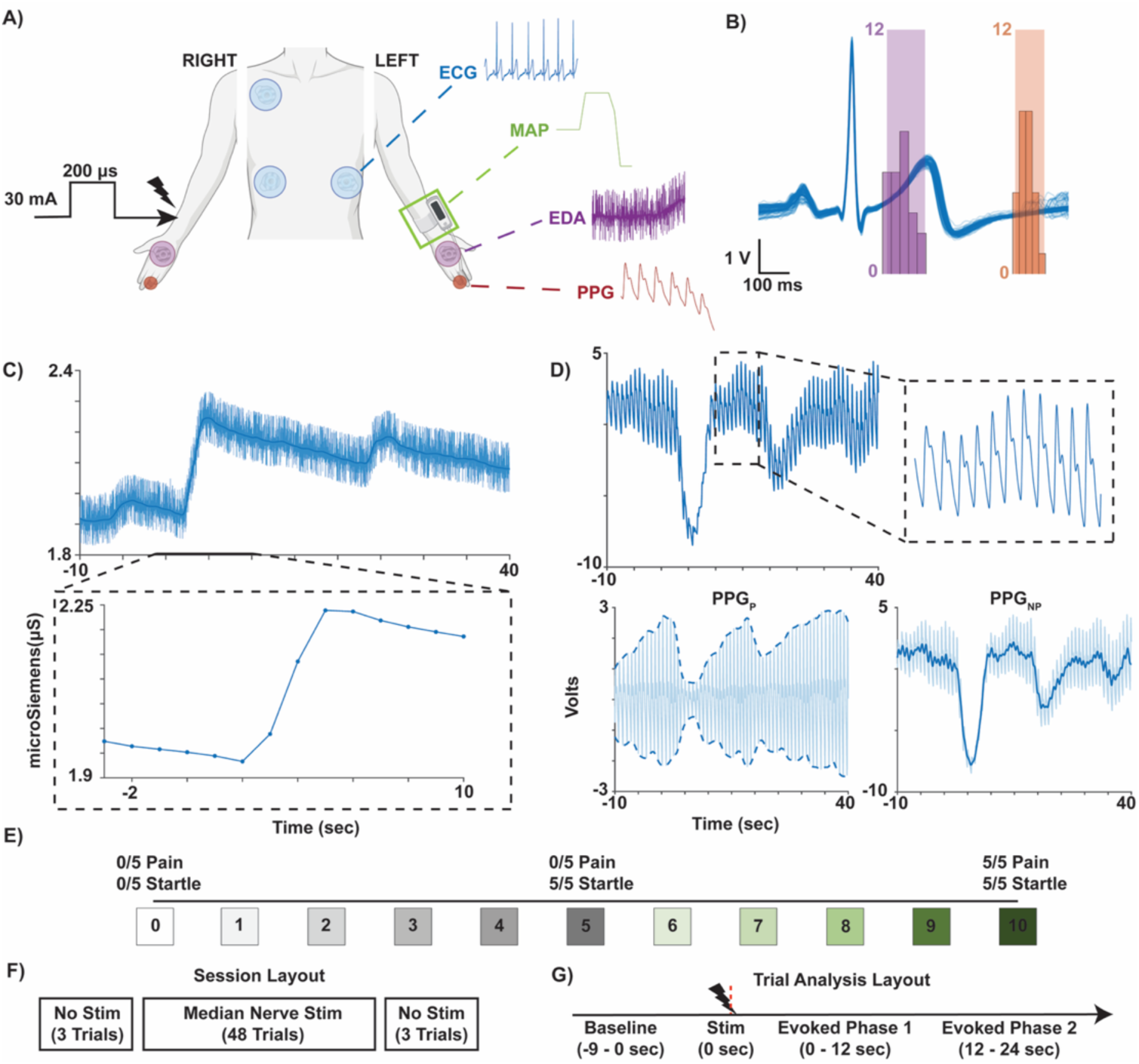
Overview Of the Experiment and Relevant Signal Preprocessing. **(A)** Schematic of the recorded signals and recording locations. **(B)** Lead II electrocardiogram (ECG) traces (blue) show when single cathodal pulse stimulation was delivered to the median nerve of the right forelimb from a representative recording. All recorded data were time-aligned to the stimulus delivery within the trial. **(C)** Raw EDA (top panel) was recorded in microSiemens (µS) and data were low-pass filtered (1 Hz cutoff, darker blue trace) and averaged in 1-second bins (bottom panel). **(D)** Raw PPG (top panel) was recorded by an infrared refractive photodetector within a sensor housing, split into pulsatile (PPG_P_, bottom left panel) and non-pulsatile (PPG_NP_, bottom right panel) components, and then averaged in 1-second bins. **(E)** Participants rated electrocutaneous median nerve stimulation using a Visual Analog Scale (VAS) (on a 250-mm line with 11 equispaced levels). **(F)** Schematic representation of the session layout. Baseline data recorded for 9 seconds before the electrocutaneous nerve stimulation were analyzed, and evoked responses were assessed in two phases following stimulation: 0 – 12 seconds (Phase 1) and 12 – 24 seconds (Phase 2). Figure panels were created in part with Biorender.

### 2.3 Median Nerve Stimulation

Stimulation electrodes were placed over the right distal median nerve approximately 1 to 2 cm proximal and medial to the radial styloid process. The electrodes were oriented 2 cm apart such that the electric field was oriented in the direction of afferent orthodromic activity within the median nerve (i.e., anode distal to cathode). During the experimental session, a constant-current stimulator (DS7R, Digitimer, Hertfordshire, UK) delivered single cathodal pulses during either systole or diastole of the cardiac cycle, as determined from a lead II electrocardiogram (ECG100C, Biopac, Goleta, CA) sampled at 1 kHz, similar to previous studies^16^. Ranges of systole and diastole, along with the distribution of stimuli delivered, are shown for a representative participant in **Figure 1B**. The current amplitude delivered by the electrode was set to 30 mA and was adjusted on a per-subject basis.

We used a discretized visual analog scale (VAS), 250 mm long with 11 equidistant levels (0 through 10) and “5” as the midpoint, to quantify the participant’s perception of the single-pulse stimulation based on how startling and/or how painful the participant felt the stimulus was. A VAS score ≤ 5 indicated the stimulus was startling but not painful, and a VAS score > 5 indicated the stimulus was startling and painful. Two participants had VAS scores greater than 5 and were unable to tolerate multiple stimulation trials. For these participants, the stimulus intensity was titrated down to tolerable amplitudes, resulting in a VAS score of 6. The VAS > 5 threshold was chosen because 5 was placed as the exact midpoint of the scale. The final stimulation amplitudes across enrolled participants align with published studies delivering median nerve stimulation in the range of 9 mA to 100 mA^28,31,49^. Only 1 of 15 (7%) participating individuals required stimulation amplitudes below 25 mA. Stimulus amplitudes and their associated VAS ratings for individual participants are provided in **Supplemental Table S1** with an average recorded VAS of 4.6 across all participants.

### 2.4 Signal Data Collection

#### 2.4.1 Electrodermal Activity

Electrodermal activity (EDA) (associated with sudomotor activity) was collected bilaterally using Ag/AgCl electrodes placed on the dorsal and ventral surfaces of the palm, streamed through a Bluetooth signal processor (Plux Signals, Lisbon, Portugal), and sampled at 1 kHz. Data (in microSiemens or µS) were low-pass filtered with a cutoff of 1 Hz using a fourth-order Butterworth filter (to remove 60-Hz noise). One thousand millisecond time windows with no overlap were used to bin the data over the time course of the trials. A visual summary of the EDA signal preprocessing during a representative trial is shown in **Figure 1C**.

#### 2.4.2 Photoplethysmography

Photoplethysmogram (PPG) signals (associated with skin vasomotor activity) were collected bilaterally using optical transducers (TSD200C) and associated preamplifiers (PPG100C) manufactured by BIOPAC (Goleta, CA, US). Each PPG sensor was placed on the finger pad of the fourth digit of each hand. Wires were taped to the base of participants’ wrists to minimize artifacts caused by wire movement. PPG signals contained correlates of underlying changes in blood volume. Signal preprocessing was performed in accordance with published studies to extract the pulsatile (PPG_P_) and nonpulsatile (PPG_NP_) blood volume indices^50,51^ (similar to the well-reported PPG-based perfusion index). The PPG_P_ blood volume index was obtained by removing high frequency signal from the raw PPG data^50,51^ using a fourth-order Butterworth Filter (1–15 Hz). The mean absolute envelope of the filtered signal was binned using 1000 millisecond non-overlapping windows. The PPG_NP_ component of the PPG signal was obtained by low-pass filtering the raw signal at 1 Hz (4th-order Butterworth Filter) and binning with a 1000-millisecond non-overlapping time window. A visual summary of the preprocessing for both the PPG_P_ and PPG_NP_ blood volume indices during a representative trial is shown in **Figure 1D**.

##### 2.4.2.1 Validation of PPG Signals

To validate the PPG signals, a reactive hyperemia study was conducted in a random subset of participants (n = 2 males and n = 2 females). A manual sphygmomanometer (Hillrom, Welch Allyn Inc., NY, USA) was inflated over the brachial artery to a measured pressure of ∼20 mmHg above the participant’s measured systolic BP to occlude blood flow, while recording PPG signals from the finger pad of the 4^th^ digit distal phalanx. The cuff remained inflated for 30 s. PPG signals were recorded for 60 s before and after inflation, in addition to the 30 s during inflation. PPG was then preprocessed accordingly to separate PPG_P_ and PPG_NP_. PPG_P_ data was normalized to full trial data according to the following formula:

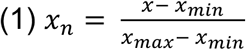

(where x_n_ is a given normalized data point; x is a given unnormalized data point; x_min_ and x_max_ are the minimum and maximum data points within the data set, respectively). PPG_NP_ data were standardized to mean and standard deviation of data collected from 10- to 50-seconds according to the following formula:

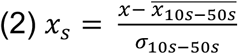

(where x_s_ is a given standardized data point; x is a given unstandardized data point; 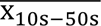 and σ_10s-50s_ are the mean and standard deviation of data points between time points 10 s and 50 s, respectively).

#### 2.4.3 Electrocardiography

A lead II electrocardiogram (ECG) recording was obtained using Ag/AgCl electrodes attached to positive, negative, and ground leads placed on participants as illustrated in **Figure 1A**. Leads were connected to an ECG100C preamplifier via an MEC110C extension cable (BIOPAC, Goleta, CA, US). Data obtained via BIOPAC preamplifiers (including PPG and ECG) were streamed via a National Instruments Data Acquisition and Control unit (Multifunction BNC I/O USB-6343, National Instruments, Hungary). ECG data were notch filtered using a fourth-order Butterworth filter. Single-pulse cathodal stimulation was randomly delivered to the right median nerve either during systole (R-wave + 150ms; purple distribution in **Figure 1B**) or diastole (R-wave + 650ms; orange distribution in **Figure 1B**) to assess possible gating of responses due to cardiac cycle timing.

#### 2.4.4 Beat-by-Beat Cardiovascular Recordings

The following beat-by-beat cardiovascular signals were recorded using the Vitalstream system (Caretaker Medical, Charlottesville, VA, US): MAP, HR, SV, and CO. A low-pressure finger sensor was placed on the middle phalanx of the left third finger to transmit data to the CareTaker wrist module. Although the device records CO and SV, thermodilution is required to validate their actual values. Thus, CO and SV reported herein only represent changes from baseline (delta or Δ). Brachial artery BP measured via digital cuff sphygmomanometer (Omron Healthcare Co. Ltd., Kyoto, Japan) before the start of the session was used to validate the Vitalstream System’s accuracy for MAP in each participant.

### 2.5 Signal Analysis

#### 2.5.1 Quantification of Evoked Responses

Data processing was conducted in MATLAB 2024a (MathWorks, Natick, MA). Signals were time-aligned where 0 seconds corresponded to the time of stimulus delivery. After preprocessing, the mean signal trace was obtained across all trials for each participant. The mean signal trace for each participant was then z-scored to its respective mean during the baseline period (t < 0). Trapezoidal approximation area under the curve (AUC) over 3-s non-overlapping time intervals was obtained from the z-scored data. Data recorded from peripheral signals were compared across ipsilateral and contralateral sites relative to the stimulation electrodes over the right median nerve.

Our pilot studies suggested multi-phasic responses in a subset of recorded signals. Therefore, the mean evoked activity for each signal was evaluated across 3 response phases: baseline (-9 to 0 s), Evoked Phase 1 (0 to 12 s), and Evoked Phase 2 (12 to 24 s). Pearson correlations were also computed for all pairs of recorded signals. For signal data exhibiting a significant change over time (i.e., a significant main effect of time in a repeated-measures ANOVA), effect sizes were calculated and ranked to identify the most sensitive signals evoked by startle.

#### 2.5.2 Assessment of Response Latency Properties

Response latency properties were also derived for each signal. Latency herein is defined as the amount of time from stimulation at t = 0 seconds to a relevant feature in the signal after crossing the 2*SD threshold. This method was used to obtain the following latency properties: onset latency (i.e., latency to response onset), peak latency (i.e., latency to maximum excursion from baseline), and termination latency (i.e., latency to response termination). For biphasic data, the latency to maximum excursion for the two relevant features were obtained.

#### 2.5.3 Assessment of Response Gating

Evoked responses to stimuli, such as startling peripheral electrical stimulation, Valsalva, or auditory startle may depend on sex or cardiac cycle phase (i.e., systole or diastole, corresponding to high or low baroreceptor load, respectively)^16,52–54^. Thus, evoked responses across signals were categorized and compared by: (i) the participant’s sex, or (ii) phase of the cardiac cycle when the stimulus was delivered (systole (R-wave + 150 ms) or diastole (R-wave + 650 ms)). As described above, the calculated AUC values over non-overlapping 3-s time windows were compared to assess the presence of sex-specific or cardiac-cycle-specific gating of SNS startle reflex signal responses.

#### 2.5.4 Assessment of Response Reliability and Differential Control

To test interdependence of peripheral non-somatomotor signals and, thus, potential differential control of SNS responses^55–59^, 8 epochs consisting of 6 of AUC timeseries EDA, PPG_P_, PPG_NP_, and MAP data were concatenated together into a Boolean vector identified by subject, trial, and time-series timepoint. Data were then analyzed using a Linear Mixed-effects model (LMM). Results are reported as standardized linear coefficient estimates (β). Data variance across all participants and trial epochs were also analyzed to assess signal consistency and reliability across time.

#### 2.5.5 Assessment of Response Probability and Predictive Performance

Preliminary studies and published literature suggest that not all participants exhibit large-magnitude sympathetic responses to single trials of startling stimuli^29,60^. To assess response probability and predictive performance, each trial was first classified as responsive (1) or non-responsive (0) within each signal (responsive trials occurred when the AUC during evoked phase 1 was greater than the mean AUC ± 2*SD at baseline). This binary variable was applied to each trial within each participant. After fitting a Generalized Linear Mixed Model (GLMM) (using the subfamily “binary”), the estimated response probability was plotted against trial number for strong responders (upper 50^th^ percentile AUC responses) vs. weak responders (lower 50^th^ percentile AUC responses).

### 2.6 Statistical Assessments

All data collected (survey and signal data) were de-identified, preprocessed, and analyzed in a single-blind design. For data included in sections 2.5.1-2.5.3, statistical analyses were performed using GraphPad Prism. A two-way repeated-measures (RM) ANOVA was used to assess the time-series modulation of the signals at two independent variable levels across subjects. For example, in section 2.5.1 where single signals within an independent variable level were analyzed, a one-way RM ANOVA was used. This statistical design was also used for analyses conducted in section 2.5.2. Note that all post-hoc analyses were either Dunnett corrected for comparison of each measure to a single control time-point (baseline period), or Tukey corrected for multiple comparisons across groups, as appropriate. For section 2.5.1, a Pearson correlation analysis was conducted to assess the covariance among all dependent variables. The ranking of relevant within-subjects time factor effect sizes from relevant RM ANOVAs was also conducted in section 2.5.1. For data in sections 2.5.4 and 2.5.5, statistical analyses were conducted in R (v. 4.4.1; R Core Team, 2024) using LMM functions from the R nlme package^61^ (v. 3.1.166) applied to the concatenated trial epoch signal data, and a GLMM from the lme4 package^62^ (v. 1.1-35) applied to the binary responsivity signal data across all trials within each participant, respectively. For section 2.5.4, signal fixed factors were analyzed for bidirectional effects (e.g., EDA ∼ PPGP + PPG_NP_ + MAP + Time). This resulted in 4 models, which were Bonferroni corrected. For section 2.5.5, overall odds ratios and standard errors (SE) across all peripheral signals were obtained as a weighted average and pooled SE, respectively, according to the following equations:

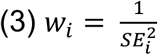

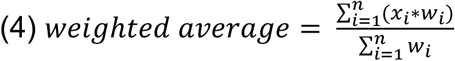

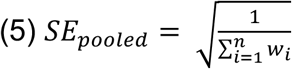

(where *w* is the weight based on the SE of the calculated log-odds *x* across each of n = 3 signals (EDA, PPG_P_, and PPG_NP_)). Bonferroni corrected post-hoc estimated marginal mean (EMM) comparisons were conducted to compare either groups or signals.

## 3. RESULTS

### 3.1 Validation Experiments for Electrocutaneous Nerve Stimulation & PPG Signals

We first confirmed that electrocutaneous nerve stimulation activates the startle reflex (similar to previous studies^8,18^) by comparing evoked signal responses with a traditional startling auditory stimulus (1-kHz, 200-ms, 105 dB)^4^ in a random subset of participants (n = 3). As expected, evoked responses using either stimulation modality demonstrated similar response profiles across the EDA, PPG_P_, and PPG_NP_ signals (**Supplemental Figure S1**). This demonstrates that electrocutaneous nerve stimulation can activate the startle reflex. We also initially characterized and validated the PPG signals to confirm that changes in signal level correspond to changes in relative blood volume (using a reactive hyperemia test in a random subset of participants (n = 4)). PPG was recorded from the 4^th^ digit, and supra-systolic pressure was applied proximally via a sphygmomanometer on the upper arm (therefore cutting off forward arterial flow through the brachial artery). PPG_P_ decreased significantly from baseline to an absolute value of 0 during the occlusion period (**Supplemental Figure S2**). In contrast, the PPG_NP_ signal showed an overshoot exceeding the initial decrease from baseline, suggestive of volume transfer from high- to low-pressure vascular compartments. These PPG signal validation results demonstrate that a decrease or increase in the recorded PPG signal corresponds to a decrease or increase in the respective blood volume.

### 3.2 Startling Electrocutaneous Stimulation Evokes a Bilateral Peripheral Startle Response

Across all participants (n=15), electrocutaneous stimulation evoked monophasic deflections relative to baseline in EDA and PPG_P_ (**Figure 2A-B**). EDA exhibited an average baseline AUC of approximately zero (0.01 ± 0.02) before stimulation (t < 0). Following stimulation at t = 0, EDA values increased to a mean AUC of 23.01 ± 6.99 with a maximum excursion from baseline occurring during the evoked phase 1 between 9 and 12 seconds (**Figure 2A – left**). After reaching maximum excursion, the stimulus-evoked EDA response remained elevated over baseline during evoked phase 2 between 12- and 24-second post-stimulation (**Figure 2A – right**). Similar, but opposite in polarity to EDA responses, PPG_P_ values began at a baseline mean AUC of 0.05 ± 0.07 and decreased to a mean maximum AUC of -9.68 ± 2.50 during evoked phase 1 between 9- and 12-second post-stimulation (**Figure 2B – left**). After reaching maximum excursion, the stimulus-evoked PPG_P_ response returned to baseline during evoked phase 2 between 12- and 24-seconds post-stimulation (**Figure 2B – right**). These data support the hypothesis that an increased sweat response and a decreased pulsatile blood volume index (indicative of skin vasoconstriction) can be reliably evoked following startling electrocutaneous stimulation.

**Figure 2.**
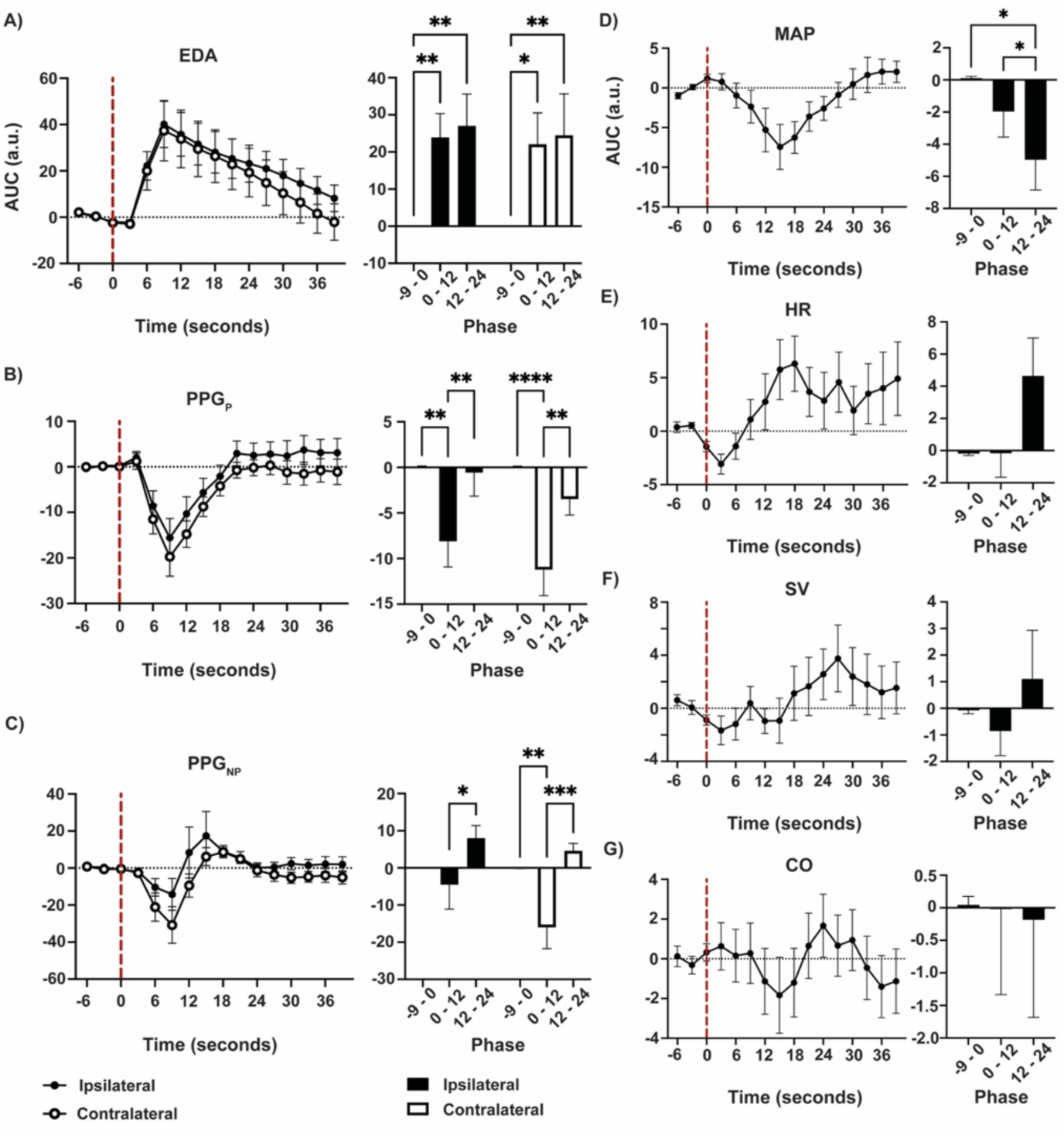
Bilateral Evoked Responses from Sudomotor, Vasomotor, and Cardiovascular Signals Following Startling Electrocutaneous Stimulation. All recorded peripheral signals (except ipsilateral PPG_NP_) demonstrated excursions from baseline following electrocutaneous stimulation **(Left Column).** There were no significant differences between peripheral signals recorded either ipsilateral or contralateral to the stimulation site. EDA data **(A)** showed a sustained monophasic increase following stimulation that persisted above baseline in both evoked phases. Pulsatile PPG (PPG_P_) **(B)** blood volume index demonstrated monophasic responses with initial decreases in signal amplitude, while nonpulsatile PPG_NP_ **(C)** signals displayed a biphasic response profile. MAP **(D)** was the only central hemodynamic signal to display a significant evoked response following stimulation. There were no significant changes in HR **(E)**, ΔStroke Volume (SV) **(F)**, or ΔCardiac Output (CO) **(G)** following stimulation. Line graphs (left subpanels) contain mean ± SEM evoked signal responses to right median nerve electrocutaneous stimulation (delivered at t = 0s; red dashed line) across time for ipsilateral (black circles) and contralateral (white circles) channels. Right subpanel bar graphs show mean AUC data ± SEM across the three phases for both ipsilateral (black bar) and contralateral (white bar) channels: Baseline (< 0 seconds), Evoked Phase 1 (0 – 12 seconds), and Evoked Phase 2 (12 – 24 seconds) (*, p < 0.05; **, p<.01; ***, p < 0.001; **** p < 0.0001).

In contrast, PPG_NP_ exhibited a biphasic response waveform (**Figure 2C – left**), associated with changes in venous-capillary blood volume. PPG_NP_ AUC values were close to zero (-0.02 ± 0.05) during the baseline period and subsequently decreased to -10.31 ± 5.44 during evoked phase 1. Similar to PPG_P_, this decrease in signal is indicative of a reduction in blood volume index. However, during evoked phase 2, PPG_NP_ AUC returned to and transiently overshot baseline values, reaching maximum excursion at 6.27 ± 2.40 (**Figure 2C – right**). While the initial decrease in PPG_NP_ aligned with observed changes in the PPG_P_ signal, the evoked phase 2 overshoot in PPG_NP_ was unexpected (potentially associated with complex skin hemodynamics; see Discussion).

Bilateral responses were observed across all peripheral signals following electrocutaneous nerve stimulation. Overall, evoked EDA (**Figure 2A**, p = 0.69), PPG_P_ (**Figure 2B**, p = 0.32), or PPG_NP_ (**Figure 2C**, p = 0.17) responses recorded either ipsilateral or contralateral to the stimulation site (i.e., the right median nerve) were not significantly different. These results support the hypothesis that evoked responses mediated by the startle reflex are associated with equal bilateral outflow, inducing changes in sweating and blood volume indices.

### 3.3 Startling Electrocutaneous Nerve Stimulation Evokes Mild Changes in Central Cardiovascular & Hemodynamic Signals

Startle reflex activation can induce central cardiovascular and hemodynamic changes following a severe startling or prolonged stimulus^12,15,17^, compared to the more mild stimulus used here. Out of the four central signals recorded (MAP, HR, ΔCO, & ΔSV), only MAP data demonstrated significant changes over time (**Figure 2D**). Mean MAP AUC decreased with time after nerve stimulation to a maximum excursion from baseline of -4.97 ± 1.88 at the end of the evoked phase 1 from 0.10 ± 0.12 in the baseline period. During evoked phase 2, mean MAP AUC returned to baseline levels.

Although there were no statistically significant main effects of time for the other central signals (**Figure 2E-2G**), notable trends emerged. Mean HR AUC **(Figure 2E)** demonstrated a trend towards a small decrease from -0.20 ± 0.12 during the baseline phase to a maximum of -0.37 ± 1.54 during the evoked phase 1. After reaching maximum excursion, the mean HR AUC trend returned to and overshot baseline values during the evoked phase 2. Neither ΔCO nor ΔSV displayed consistent or notable trends (**Figure 2F-G**). For this reason, ΔCO and ΔSV were not considered for further analyses. Overall, in contrast to the array of robust peripheral signal responses, evoked central responses were generally mild and limited to MAP.

### 3.4 Comparison of Evoked Response Latencies

Regarding the temporal dynamics of responses, the onset latency for MAP was significantly longer compared to the peripheral signals (**Figure 3A**). MAP peak latency generally coincided with the second maximum excursion of the biphasic PPG_NP_ response (p = 0.60), with the peak latency for both signals being significantly longer than the remaining peripheral signals (**Figure 3B**). Lastly, MAP and PPG_NP_ took the longest to terminate, with MAP being significantly longer than PPG_P_ (**Figure 3C**). Overall, these results outline the temporal structure of SNS responses following startle.

**Figure 3.**
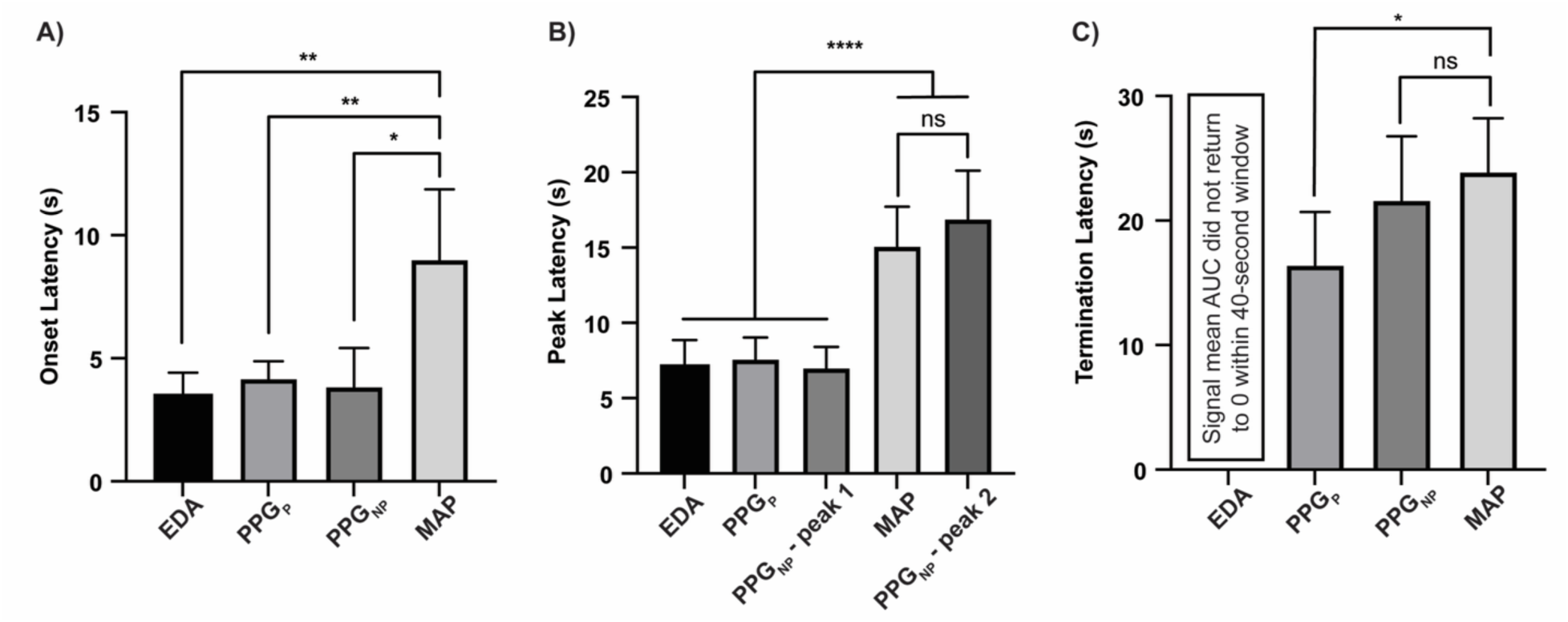
Temporal Response Structure of Evoked Responses Following Startle. Onset latency and peak latency for peripheral signal responses were typically ∼50% faster compared to MAP responses (**A** and **B**, respectively). MAP responses displayed a peak latency (**B**) coincident with the second PPG_NP_ peak within evoked phase 2 (between 14 and 18 seconds). (**C**) EDA did not return to baseline during the 40 s post-stimulation window, while both PPG_NP_ and the MAP generally terminated together. PPG_P_ responses terminated significantly faster than MAP responses. Data are presented as mean ± SEM (*, p < 0.05; **, p<.01; ****, p < 0.0001).

### 3.5 The ‘SNS Startle Signature’ and Comparison of Evoked Responses for Peripheral and Central Signals

Historically, the human startle reflex has been primarily characterized through its coordinated and highly patterned somatomotor expression^18–21^. Comparatively fewer studies have characterized the accompanying autonomic and sympathetic responses^12,15,38,63–65^, and even fewer have simultaneously assessed multiple sympathetic effector systems. Here we provide evidence for a coordinated, patterned, and multimodal SNS startle signature (**Figure 4A)**. The SNS startle signature is comprised of the simultaneously activated EDA, PPG_NP_, PPG_P_, and MAP signals (i.e., only signals with statistically significant changes). Furthermore, the SNS startle signature exhibits a highly repeatable and patterned response with low variance across subjects. Overall, this result complements several important prior works demonstrating a coordinated and highly patterned somatomotor response following startle and extends these findings to SNS-linked signals.

**Figure 4.**
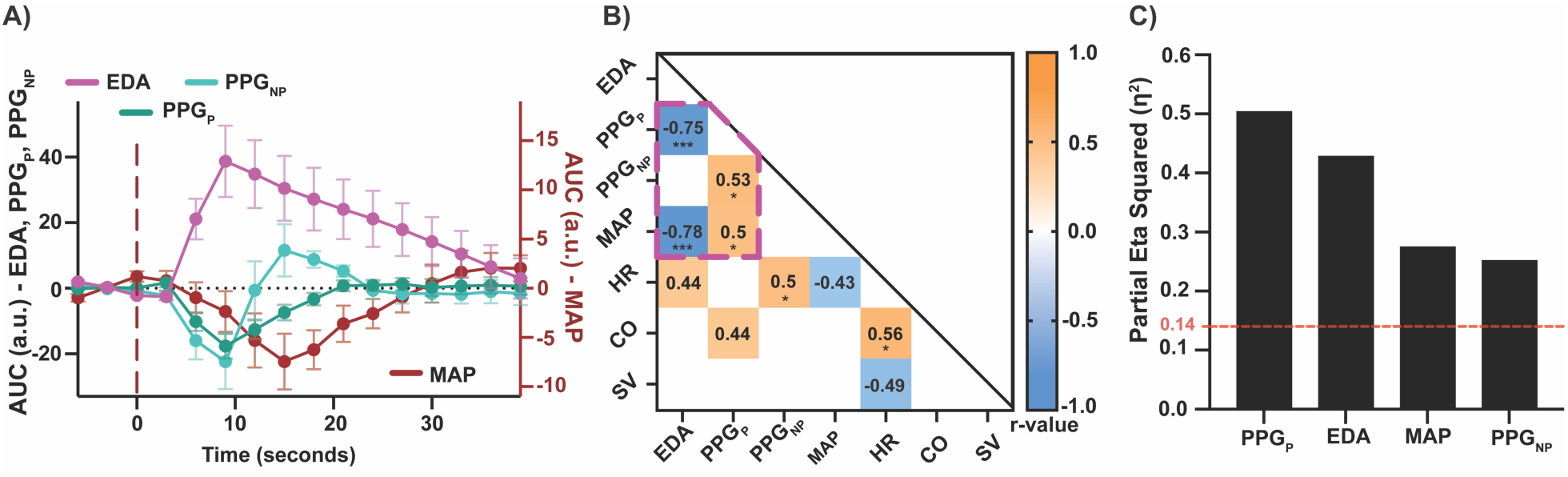
The SNS Startle Signature and Comparison of Evoked Responses for Peripheral and Central Signals. **(A)** The SNS startle signature was comprised of a highly coordinated and repeatable set of response profiles across the EDA, PPG_NP_, PPG_P_, and MAP signals (data are presented as mean ± SEM). **(B)** Overall, multiple peripheral and central signal responses were significantly correlated. Pairs of signals within the dashed pink outline denote significant correlations that also displayed a significant main effect of time (see Figure 2). Increases in sweat response (EDA) correlated with decreases in pulsatile blood volume (PPG_P_). MAP also had an inverse correlation with EDA and a weak positive correlation with PPG_P_. **(C)** Effect size ranking (partial eta squared (η^2^)) is presented in descending order for signals demonstrating significant main effects of time (i.e., EDA, PPG_P_, PPG_NP_, and MAP). The red horizontal line indicates the threshold for a large effect size based on partial-η^2^ *^66^*. Three of the four signals showing large effect sizes were peripheral signals. The partial-η^2^ for MAP was similar compared to PPG_NP_. Taken together, these data demonstrate the existence of a multimodal SNS startle signature and further characterize its overall response structure (*, p < 0.05; ***, p < 0.001).

Regarding response patterns, Pearson correlation analysis revealed that both peripheral and central signals significantly correlated with each other following startle (**Figure 4B**). Furthermore, peripheral evoked responses (EDA & PPG) were significantly larger compared to central evoked responses (MAP & HR) (**Supplemental Figure S4**). Lastly, using effect size, peripheral signals outperformed central signals approximately 2:1, again indicating greater overall evoked changes for peripheral signals (**Figure 4C**).

### 3.6 Neither Biological Sex nor Cardiac Phase Affects Evoked Peripheral and Central Signal Responses

Previous studies have reported a dependence of startle responses on subject sex (male vs. female) and cardiac phase (i.e., during systole or diastole)^16,52–54^. To assess these potential response dependencies, both male and female participants were separately assessed, and startling electrocutaneous nerve stimulation was delivered during either systole or diastole of the cardiac cycle based on a live-streamed lead II ECG (with randomized but equal allocation across 48 trials; **Figure 1B**). Analysis revealed that neither sex (**Supplemental Figure S5A**) nor cardiac cycle (**Supplemental Figure S5B**) affected startle-evoked EDA, PPG_P_, PPG_NP_, MAP, and HR responses. Only mild trends were observed, where startle-evoked responses tended to be larger when the participant was female or when the stimulus was delivered during systole. While these trends do, in part, align with previously published studies demonstrating larger startle responses in females^54^, the data presented herein suggest that startle reflex activation following electrocutaneous stimulation was not significantly affected by sex or cardiac phase.

### 3.7 Evoked Responses Following Startling Electrocutaneous Stimulation In Part Exhibit Differential Control of Sympathetic Outflow

Several studies have recorded SNS-linked responses following startle reflex activation, but relatively few have assessed these responses through the established framework of differential control of sympathetic outflow^55–59^. Similar to previous studies, we define differential control as evoked responses that either do not respond together or do not respond with equal weight^55,56^. Possible changes (β) in EDA (**Figure 5A)**, PPG_P_ (**Figure 5B)**, PPG_NP_ (**Figure 5C)**, and MAP (**Figure D)** are shown as a function of unit changes across the remaining reference signals. Our results show that 8 of the possible 12 coordinated responses between signals were statistically significant and were not necessarily of equal weight β. For example, unit changes in PPG_P_ were predictive of changes in EDA (**Figure 5A**), PPG_NP_ (**Figure 5C**), and MAP (**Figure 5D**). However, a unit change in EDA was only predictive of a mild change in PPG_P_ (**Figure 5B**). Furthermore, evoked changes were also consistent and repeatable throughout testing sessions, as signals from each participant did not significantly deviate from the global average, indicating high response reproducibility and reliability (**Figure 5E-H**). An overview of the coordinated responses across signals is shown in **Figure 5I**. Taken together, these data suggest that startle-evoked responses are in part under differential control of sympathetic outflow^55–59^.

**Figure 5.**
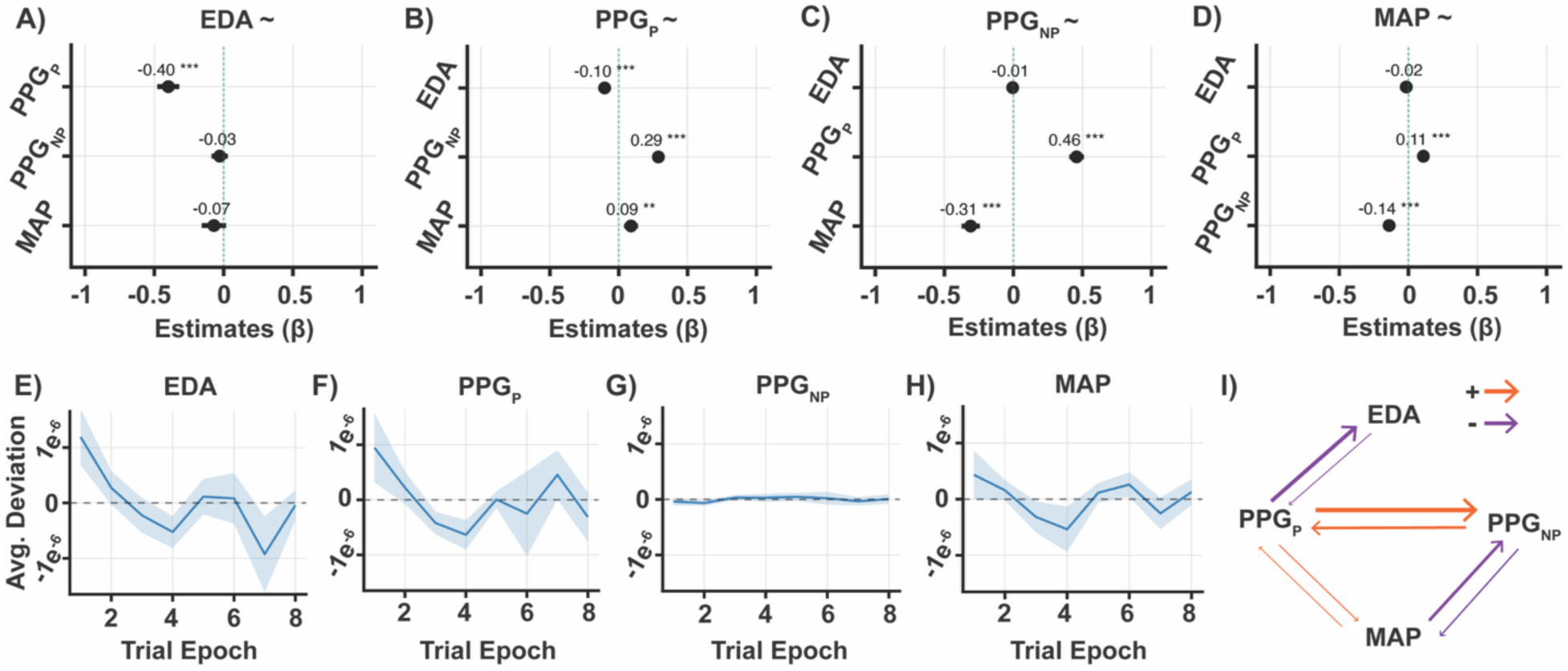
SNS Startle Responses Are In Part Under Differential Control of Sympathetic Outflow. (**A-D**) Forest plots of fixed factor predictor effects across signals. The majority of signals demonstrated coordinated responses. However, subsets of responses demonstrated differential control. For example, unit changes in PPG_NP_ and MAP did not predict changes in EDA (**A**). Furthermore, unit changes in EDA did not predict changes PPG_NP_ (**C**) or MAP (**D**). (**E-H**) Responses were reproducibile and reliabile across the testing population with low magnitude deviations across trials. (**I**) Summary overview of coordinated responses across signals. Arrows denote the existence and directionality of a given response relationship (orange: coordinated positive response; purple: coordinated negative response). (**A-D**) Data are presented as linear coefficient effect-size, β ± CI; (**E-H**) Data are presented as mean Empirical Bayes estimates ± SE (*, p < 0.0125; **, p<0.0025; ***, p < 0.00025).

### 3.8 Responder Discrimination and Predictive Performance for SNS Startle Responses

Ideal diagnostic tools characteristically display attributes such as reliable responder discrimination and robust predictive performance. To determine the responder discrimination for startle-evoked SNS responses, binary classification of responsive and non-responsive trials was assessed across all participants, and the probability of observing a response was next quantified. Strong responders across all peripheral signals demonstrated a significantly larger composite odds ratio (OR) compared to weak responders (OR = 4.52, 95% confidence interval (CI) = [3.20, 6.40]; OR = 1.23, 95% CI = [0.92, 1.65], respectively) (**Figure 6A**). The 95% CI of the weak responder group crossed the OR = 1 threshold, indicating that startle-evoked SNS responses for this group were not significantly different from chance levels. Together, these findings demonstrate clear responder discrimination and suggest that startle-evoked SNS responses may have potential diagnostic utility for identifying SNS dysfunction in clinical populations.

**Figure 6.**
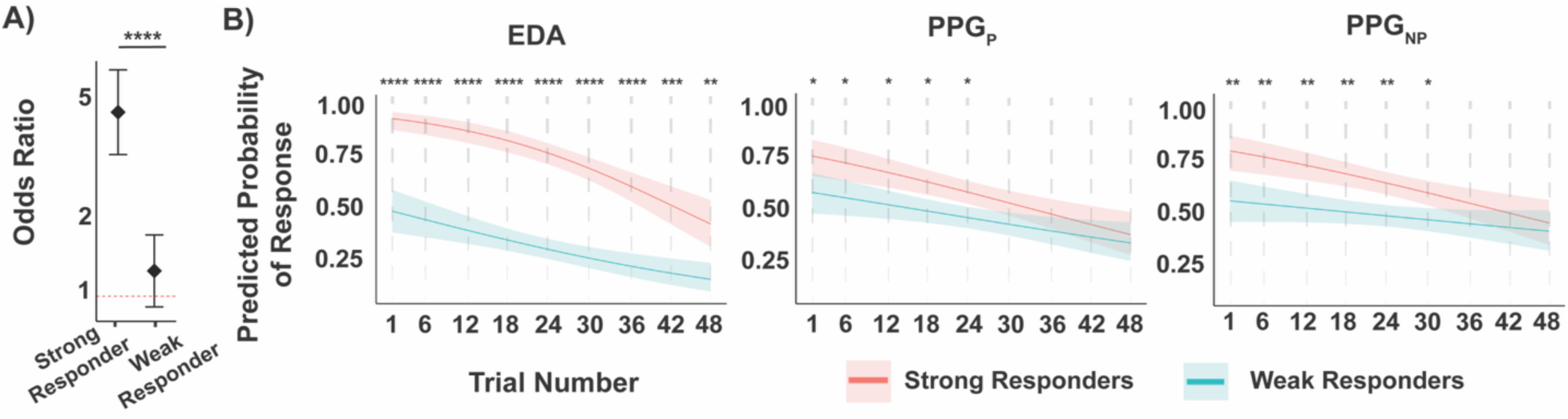
SNS Startle Responses Demonstrate Robust Responder Discrimination and Predictive Performance. (**A**) Odds ratio analysis demonstrates that strong responders were ∼3.5x more likely to display SNS startle responses compared to weak responders. (**B**) Plots from left to right display the predictive performance of each peripheral signal (EDA, PPG_P_, and PPG_NP_) for strong and weak responders across all trials of the recording session. EDA maintained a significant difference in response probability for strong and weak responders across all trials. PPG_P_ and PPG_NP_ lost significant predictive performance by trial 36 (p = 0.137) and 30 (p = 0.077), respectively. Data are presented here as odds ratio with ± 95% CI (**A**) or mean ± SEM (**B**) (*, p < 0.05; **, p<0.01; ***, p < 0.001; **** p < 0.0001).

We next assessed the response probability and general response habituation across signals and responder types. Overall, strong responders were more likely to consistently display a startle-evoked response compared to weak responders (**Figure 6B**). For EDA, the response probability for strong responders was significantly higher compared to weak responders across all trials (**Figure 6B, left panel**). The response probability for PPG_P_ (**Figure 6B, middle panel**) and PPG_NP_ (**Figure 6B, right panel)** was initially significantly higher for strong responders, but this effect was lost by trial 30 (p = 0.077) and trial 36 (p = 0.137) respectively, suggestive of diminishing predictive performance possibly from system adaptation. Overall, these results demonstrate the responder characteristics and predictive performance of SNS signals following startle.

## 4. DISCUSSION

The present study demonstrates that the startle reflex triggered by electrocutaneous nerve stimulation evokes significant changes across an array of peripheral and central signals (i.e., EDA, PPG_P_, PPG_NP_, and MAP), activating a reliable and highly structured SNS startle signature. Among healthy participants, startling nerve stimulation evoked robust changes in noninvasive peripheral sweat and vascular signals, as well as more modest changes in central signals. Peripherally recorded EDA, PPG_P_, and PPG_NP_ signals also demonstrated bilateral responses. For the central signals (MAP, HR, ΔCO, and ΔSV), only MAP showed a statistically significant response. Further analysis revealed that peripheral signals were more sensitive and more correlated with one another compared to central signals. Additionally, further analyses indicated that SNS evoked responses, although partially correlated, were in part under differential control of sympathetic outflow^55–59^. Lastly, analysis of responder discrimination and predictive performance demonstrated that startle-triggered SNS signals are a potentially viable tool for assessing ANS dysfunction in future studies.

### 4.1 Electrocutaneous Nerve Stimulation Activates the Startle Reflex and Robust Modulation of SNS Signals

Several previous studies have used auditory stimuli for evaluating both peripheral and central biomarkers of startle reflex activation^12,17,63,65^. In addition, electrocutaneous nerve stimulation has also been used to activate the startle reflex (e.g., assessed using sympathetic skin responses in patients with SCI^9,26,28,49^ and diabetes^27,31^). In the present study, EDA increased to a maximum excursion between 6 and 9 seconds after delivery of startling electrocutaneous median nerve stimulation and subsequently declined (**Figure 2A & 3**). An increase in EDA, coinciding with cutaneous vasoconstriction (i.e., PPG_P_ and PPG_NP_; **Figure 2**), is in agreement with previous studies assessing startle reflex and orienting reflex responses^63,64^. A similar monophasic change in EDA has been described in previous startle reflex studies using healthy participants^64,67,68^. However, it should be noted that studies in subjects with paralysis where EDA recording sites may be affected by the injury, startle-evoked electrodermal responses may exhibit more than one phase^26,28^.

In addition, our results indicate that cutaneous vasomotor responses evoked by startling electrocutaneous nerve stimuli may not have the same morphology and response profile compared to EDA. Cutaneous vasoconstriction occurred following stimulation, as indicated by a decrease in the pulsatile component of the PPG (PPG_P_), generally associated with an index of arterial blood volume (**Figure 2B**). In addition, stimulation also triggered a biphasic response profile in the non-pulsatile blood volume index of the PPG (PPG_NP_), generally associated with an index of venous-capillary blood volume (**Figure 2C**). Arteriovenous reflexes and venodynamics have been interrogated with pharmacologic interventions, as well as other invasive or extracorporeal cardiopulmonary preparations^50,51,69–73^. However, to our knowledge, this is the first characterization of compartment-specific skin hemodynamic responses to startle.

The startle-evoked change in PPG_P_ mirrors peripheral vasoconstriction responses reported in previous studies^63^ and is in agreement with sympathetic outflow driving vasoconstriction in response to a startling, threatening, or dangerous stimulus. The biphasic response observed in the non-pulsatile PPG component, however, may indicate a complex interplay between the fluid and structural dynamics of the underlying tissue. Overall, PPG_NP_ is generally correlated with an index of total blood volume in the tissue underlying the sensor, most closely associated with the venous-capillary compartment^50,51^. Because PPG_NP_ is calculated as a low-pass filtered (1-Hz cutoff) and binned signal, the change in the pulsatile PPG_P_ signal does not influence the PPG_NP_ signal. As such, the biphasic response observed in the PPG_NP_ signal may be evidence of an initial vasoconstrictive event primarily in the venous vascular compartment, followed by a volumetric shift from the capillary compartment that drives volume expansion and elastic damping from the now stiffened venous compartment. In this context, volume rescue is likely due to the transient contraction and rapid relaxation of vascular smooth muscle due to the brevity of the startle stimulus delivered. We hypothesize that the volume rescue reflected in the PPG_NP_ signal following a prolonged vasoconstrictive startle event would likely exceed initial volume reductions (e.g., as seen during the reactive hyperemia validation experiment (**Supplemental Figure S2**), via autoregulatory-driven vasodilation from metabolites within the metabolically challenged tissue). The finding that the latency of the second maximum excursion in PPG_NP_ occurs after PPG_P_ signal has started to return to baseline (indicating cessation of vasoconstriction and reconstitution of arterial volume) further supports the interpretation that volume reconstitution from the capillary bed and arterial pressure may be the driver of the elastic dampening and volume overshoot in the venous compartment evidenced by the secondary PPG_NP_ signal increase (**Figure 2C**), explaining the biphasic PPG_NP_ response.

In contrast to the recorded peripheral signals, central signal responses (e.g., MAP and HR) were not entirely in agreement with previous reports^15–17^. Here, we observed that electrocutaneous nerve stimulation induced a monotonic decrease in MAP with no significant effect on HR. Decreases in BP beyond 10 seconds after a startling stimulus delivery have been reported previously. However, these effects typically followed an initial increase in BP during earlier phases of the evoked response^15,17^. Additionally, we did not observe significant HR changes as previously reported following startling auditory stimuli^17^. Validation experiments comparing evoked EDA, PPG_P_, and PPG_NP_ responses to either electrocutaneous or auditory startle stimuli show that, although different in amplitude, peripheral sweat and vasomotor responses still demonstrated similar response morphologies regardless of stimulus type (**Supplemental Figure S1**). We hypothesize that a prolonged or more severe startling stimulus would drive additional significant changes in central signals. Overall, these results demonstrate a reproducible SNS startle signature across multiple signals in response to a brief and rapid electrocutaneous startle stimulus.

### 4.2 Startle-triggered SNS Responses In Part Exhibit Differential Control of Sympathetic Outflow

SNS outflow responses to startling stimuli or stressors involve multiple systems, including the well-studied caudal pontine reticular nucleus^3,74,75^, the central autonomic network^76^, and sympathetic control areas in the brainstem^3,77^. Additionally, tactile mediated startle (e.g., electrical stimulation of the median nerve) has also been reported to correlate with increased activity in the posterior parietal and somatosensory cortex^78,79^. Overall, sudomotor activity is primarily mediated through cholinergic pathways involving the posterior hypothalamus and medullary reticular nuclei^80,81^, whereas vasomotor control, including arterial and venous constriction, is largely governed by pathways involving subcortical nuclei^13^. In summary, several cortical and subcortical regions mediate distributed autonomic responses. However, the presence of shared cortical and subcortical circuitry does not necessarily imply uniform or coupled SNS output across effector systems^55–59^, raising the possibility that startle-evoked SNS responses may be subject to differential control.

Our results support this interpretation, revealing significant correlations among specific startle-evoked SNS signals, most prominently between electrodermal activity (EDA) and photoplethysmographic (PPG) measures (**Figure 4B**), while also demonstrating clear dissociations across effector systems (**Figure 5A-D**). Overall, there were complex and reliable coordinated evoked response patterns across signals (**Figure 5E-I**). These findings suggest that startle-evoked SNS responses carry correlative but non-redundant information, consistent with partial independent control of distinct sympathetic effector pathways^55–59^. These results align with longstanding evidence that sympathetic outflow is organized in a target-specific and differentiated manner, even when driven by common central inputs. For example, differential regulation of vasomotor responses have been reported across glabrous and non-glabrous skin regions, further underscoring the specificity of sympathetic patterning^58^. Furthermore, previous work has also demonstrated that different classes of stressors, such as physical versus cognitive challenges, can evoke divergent patterns of sympathetic nerve activity across muscle and skin, despite shared engagement of brainstem and hypothalamic control regions^57^. Together, these observations support the interpretation that startle engages a distributed sympathetic control architecture in which overlapping central circuitry gives rise to differential effector-specific outflow, rather than a single globally scaled autonomic response.

### 4.3 Startle-Evoked SNS Responses Demonstrate High Predictive Performance with Potential Applications as A Diagnostic Tool

Previous studies have shown that the SNS signals assessed here are associated with startle reflex activation^12,63,64^. Our study extends these findings, demonstrating that startle-evoked SNS responses are associated with meaningful responder stratification and robust response probabilities (**Figure 6**). Odds ratio analysis demonstrated that strong responders were ∼3.5x more likely to display SNS responses following startle compared to weak responders. Importantly, signal-specific analyses revealed that while EDA responses maintained predictive performance across the recording session, cutaneous vascular measures showed gradual reductions, underscoring both shared and effector-specific dynamics following startle. Together, these findings suggest that the SNS startle signature may provide positive and negative predictive information when responses are robust, while also revealing physiologically meaningful differences in response profiles across sympathetic effector systems.

The SNS startle signature may also have potential applications as a diagnostic tool for assessing ANS dysfunction in future studies. As a comparative example, the Hoffmann (H-) reflex, one of the well-established reflex-based diagnostic tools, has long served as a reliable probe of spinal sensorimotor function (**Figure 7**). Like the H-reflex, startle activates a well-defined reflex arc. The SNS startle signature extends this framework by interrogating distributed autonomic reflex effector systems, compared to a single sensorimotor pathway alone. Whereas traditional diagnostic reflexes typically yield a unitary readout (e.g., latency or amplitude), the SNS startle signature provides a reliable multidimensional profile encompassing sudomotor, vasomotor, and cardiovascular responses. This broader physiological coverage may be particularly advantageous for detecting early or partial autonomic dysfunction, where impairments may emerge in one effector system before others. Moreover, the use of noninvasive stimulation and noninvasive recordings makes the SNS startle signature possibly well suited for repeated assessments, longitudinal monitoring, and deployment in settings where conventional autonomic testing may be impractical or difficult. Looking forward, systematic evaluation of the SNS startle signature in larger clinical cohorts may enable the development of a scalable reflex-based diagnostic tool for characterizing SNS function across a wide range of neurological, cardiovascular, and ANS disorders.

**Figure 7.**
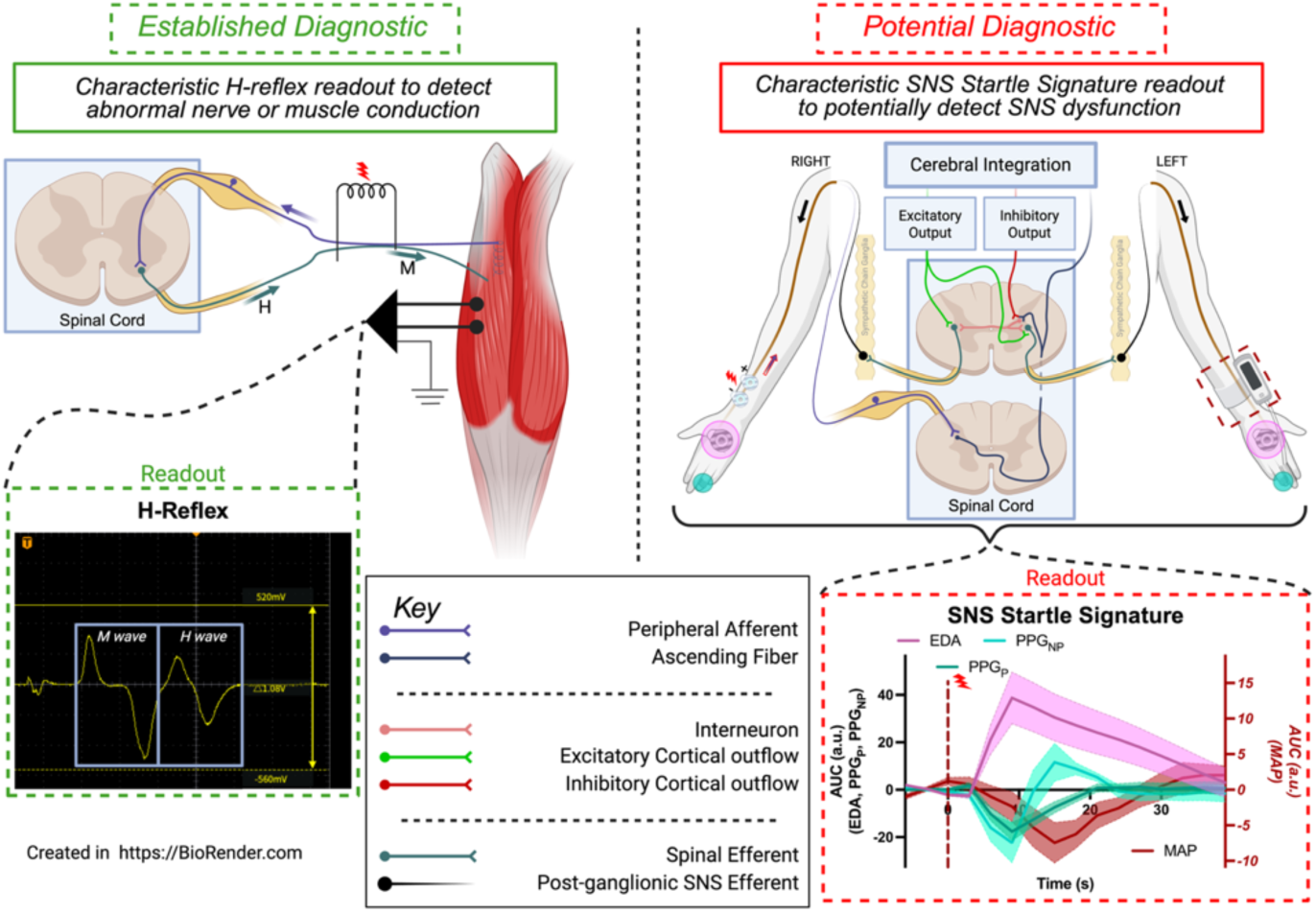
Graphical Overviews Comparing the Established H-Reflex (Left) With the SNS Startle Signature (Right). The H-reflex is a well-established reflex arc with well-characterized features that has been extensively leveraged as a diagnostic tool for assessing peripheral nerve disease, myelopathy, and spasticity (**left**). The SNS startle signature (**right-lower panel**) is a potential new diagnostic tool with possible applications for detecting SNS dysfunction (AUC = Area-under-the-curve; a.u. = arbitrary units; EDA = electrodermal activity; PPG_P_ = PPG pulsatile envelope; PPG_NP_ = PPG non-pulsatile component; MAP = mean arterial pressure). Figure panels were created in part with Biorender.

## Acknowledgements

The authors would like to thank Dr. Vivek Kanumuri, MD (University of Miami Miller School of Medicine Department of Otolaryngology, The Miami Project to Cure Paralysis, Department of Biomedical Engineering) for his contributions to the manuscript and assistance with the auditory startle experiments.

## Author Contributions

Conceptualization: RVM, ERA, GJF, DWM, & PDG. Data collection, curation, and formal analysis: RVM, ERA. Writing – Initial Draft: RVM, ERA. Writing – Editing and Revision: RVM, ERA, GJF, DWM, & PDG.

## Data Availability

All data necessary to evaluate the results presented are available throughout the paper and supplemental materials. Curated data necessary to recreate figures has been additionally made available.

## Supplemental Tables & Figures

**Supplemental Table S1.** Summary of stimulus amplitudes and associated rating of “startle” along the visual analog scale (VAS) outlined in Figure 1.

| Participant | S01 | S02 | S03 | S04 | S05 | S06 | S07 | S08 | S09 | S10 | S11 | S12 | S13 | S14 | S15 |
| --- | --- | --- | --- | --- | --- | --- | --- | --- | --- | --- | --- | --- | --- | --- | --- |
| Stimulus Amplitude (mA) | 30 | 30 | 30 | 30 | 30 | 30 | 30 | 30 | 30 | 25 | 30 | 30 | 30 | 30 | 10 |
| Stimulus Rating | 5 | 3 | 3 | 5 | 5 | 4 | 6 | 5 | 5 | 6 | 5 | 4 | 3 | 5 | 6 |

**Supplemental Figure S1.**
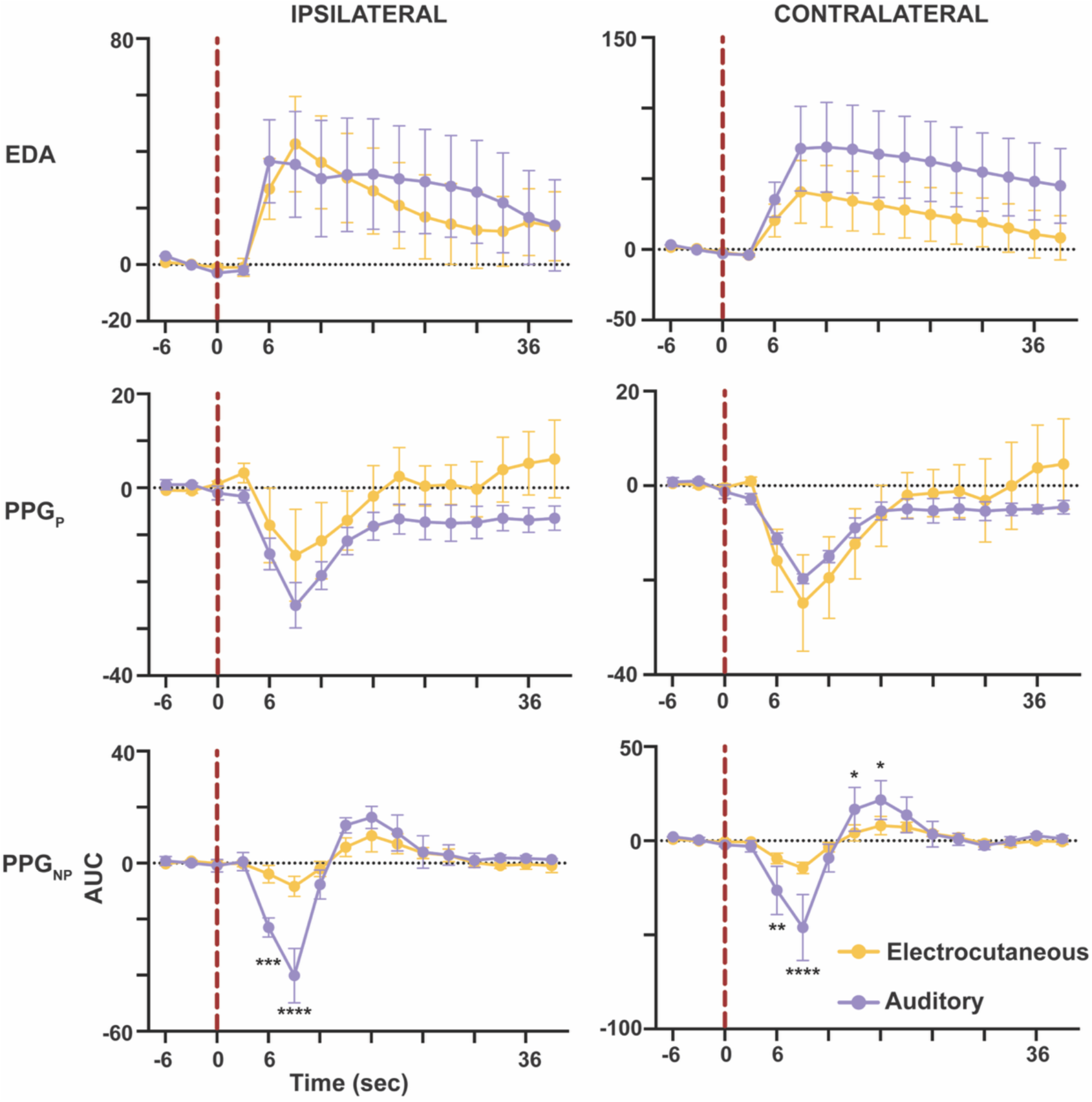
Comparison Of Evoked Peripheral Responses To Either Unilateral Electrocutaneous Or Binaural Auditory Stimulation. To in part validate the startling electrocutaneous stimulation, we compared evoked responses during either electrocutaneous stimulation or auditory stimulation within a subpopulation of the study cohort (n = 3). Auditory startle stimulation was comprised of a traditional 200 millisecond 1kHz 105 dB pure tone^4^. Aside from modest differences in PPG_NP_ amplitude, peripheral sweat and cutaneous vasomotor responses demonstrated similar response morphologies following startling electrocutaneous or auditory stimulation overall. Red dashed line = time of stimulation (acoustic startle application or electrocutaneous stimulation). Data are presented as mean ± SEM (*, p < 0.05; **, p<.01; ***, p < 0.001; **** p < 0.0001).

**Supplemental Figure S2.**
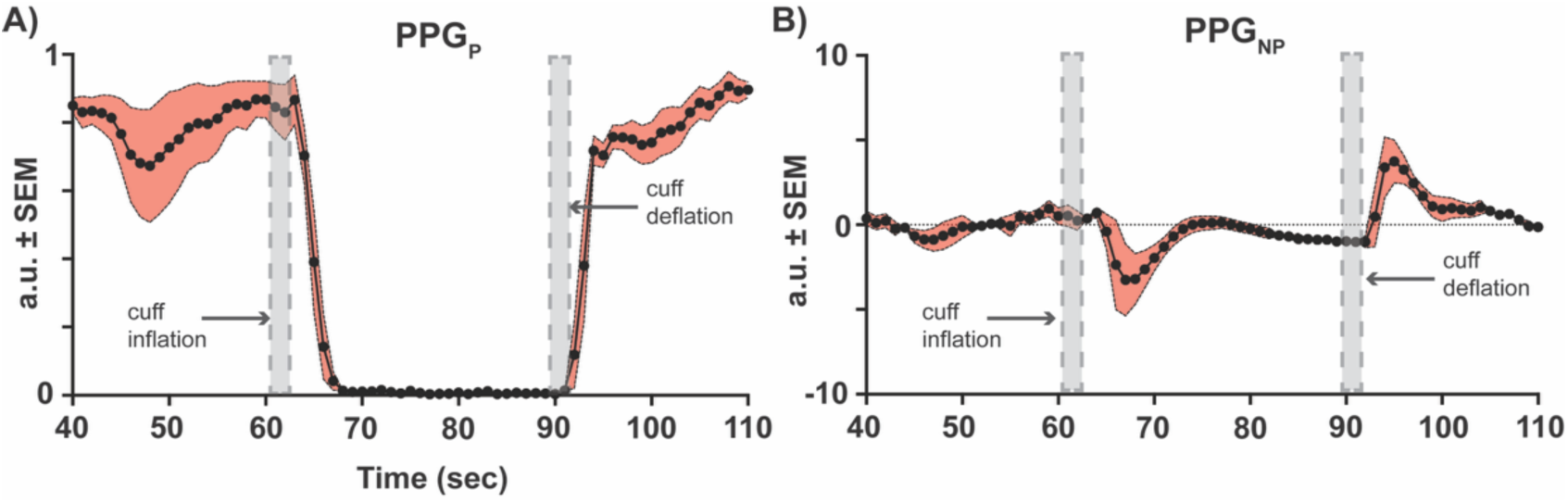
PPG Signal Validation. To validate the PPG signals, a reactive hyperemia study was conducted in a subset of participants (n = 4). Please see the *Validation of PPG Signals* in the Methods for more details. PPG signals were recorded from the hand (i.e., the finger pad of the 4^th^ digit distal phalanx) during proximal application of suprasystolic pressure over the brachial artery using an occlusion cuff. (**A**) PPG_P_ for all participants reduced to 0 during inflation of the brachial cuff. Immediately following cuff deflation (i.e., removal of suprasystolic occlusive pressure), PPG_P_ returned to baseline values. Because systolic blood pressure is defined as the maximum arterial wall pressure produced during the cardiac cycle, suprasystolic pressure was sufficient to occlude forward arterial flow to the distal PPG sensor. Occlusion of forward blood flow coincident with 0 signal amplitude in the PPG_P_ signal supports the hypothesis that decreases in PPG_P_ are correlated with decreases in underlying arterial blood volume. **(B**) PPG_NP_ initially decreased during inflation of the brachial cuff. However, during the full occlusion phase, PPG_NP_ recovered to near baseline levels. Due to the applied proximal suprasystolic occlusive pressure, no volume can return via the venous system, and importantly no volume is able to move forward in the arterial system. Therefore, volume rescue overall during the occlusion period is only possible with passive movement of existing peripheral volume from high- to low- pressure vasculature (i.e. from arterial and capillary compartments to venous and more distal capillary compartments, via arterio-venous anastomoses and venous-capillary coupling). This is further supported by the initial PPG_NP_ overshoot post-occlusion. These results support the hypothesis that following proximal suprasystolic occlusive pressure, PPG_NP_ is most influenced by volume within the venous-capillary space. Data are presented as mean ± SEM.

**Supplemental Figure S3.**
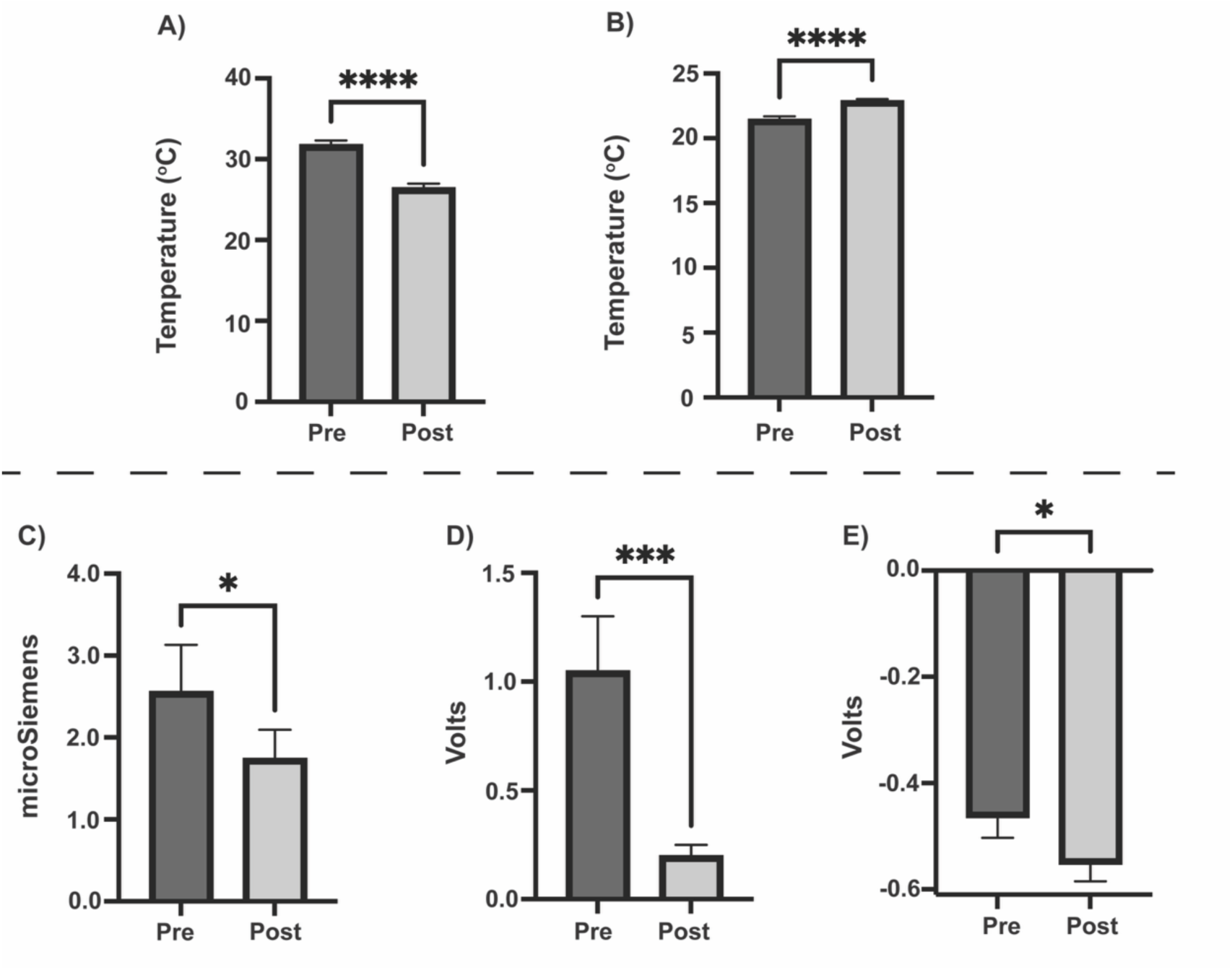
Overview Of Limb Temperature, Room Temperature, And Effects on Baseline Peripheral Signals. Summary of participant limb temperature (**A**) and ambient room temperature (**B**), as well as raw baseline EDA (**C**), PPG_P_ (**D**), and PPG_NP_ (**E**) values recorded during 3 pre- and 3 post-session trials. Participant peripheral limb temperature measured at the palms significantly decreased (**A**), while the ambient room temperature significantly increased (**B**). Raw baseline EDA (**C**), PPG_P_ (**D**), and PPG_NP_ (**E**) values significantly decreased from pre- to post- session. These results support the hypothesis that peripheral limb temperature decreases can lead to decreases in vasomotor and sudomotor signals, similar to previous studies^43,44^. Overall, decreasing baseline trends were expected in peripheral hemodynamic signals and were controlled for in analyses included in the main manuscript text. Data are presented as mean ± SEM (*, p < 0.05; ***, p < 0.001; **** p < 0.0001).

**Supplemental Figure S4.**
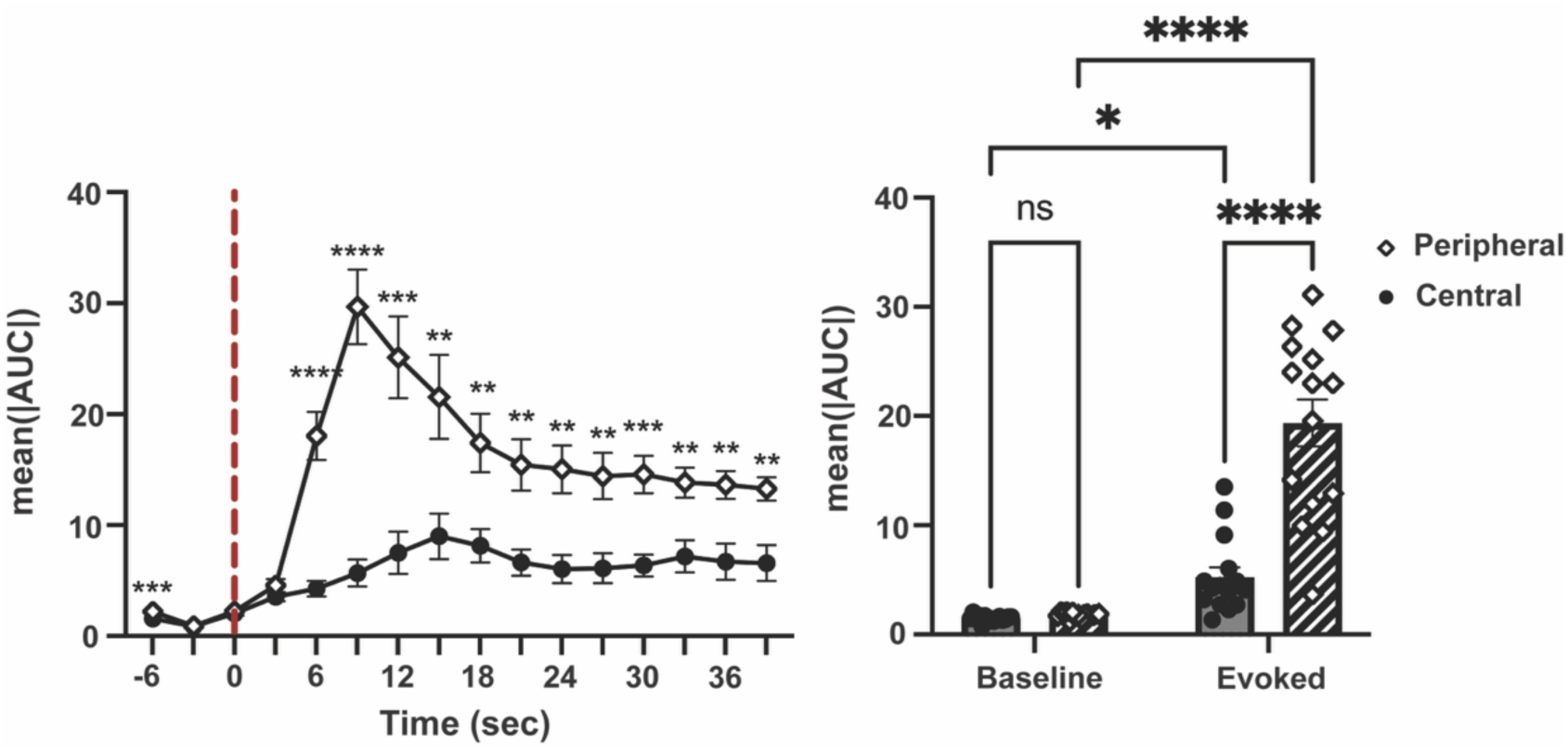
Comparison Of All Evoked Peripheral and Central Signals. **(Left panel**) Aggregate timeseries data for the evoked peripheral and central signals (central signals = MAP and HR; ΔSV and ΔCO are excluded, as they did not exhibit trends or significant changes evoked by stimulation; please see Figure 2 for more details). Peripheral signals demonstrated a greater evoked response across time compared to central signals following electrocutaneous stimulation (red dashed line = time of stimulation). (**Right panel**) There were no differences in baseline values for peripheral and central signals prior to stimulation (Baseline). Evoked response magnitudes for peripheral signals were significantly higher compared to central signals post-stimulation (Evoked). Data are presented as mean ± SEM (*, p < 0.05; **, p<.01; ***, p < 0.001; **** p < 0.0001).

**Supplemental Figure S5.**
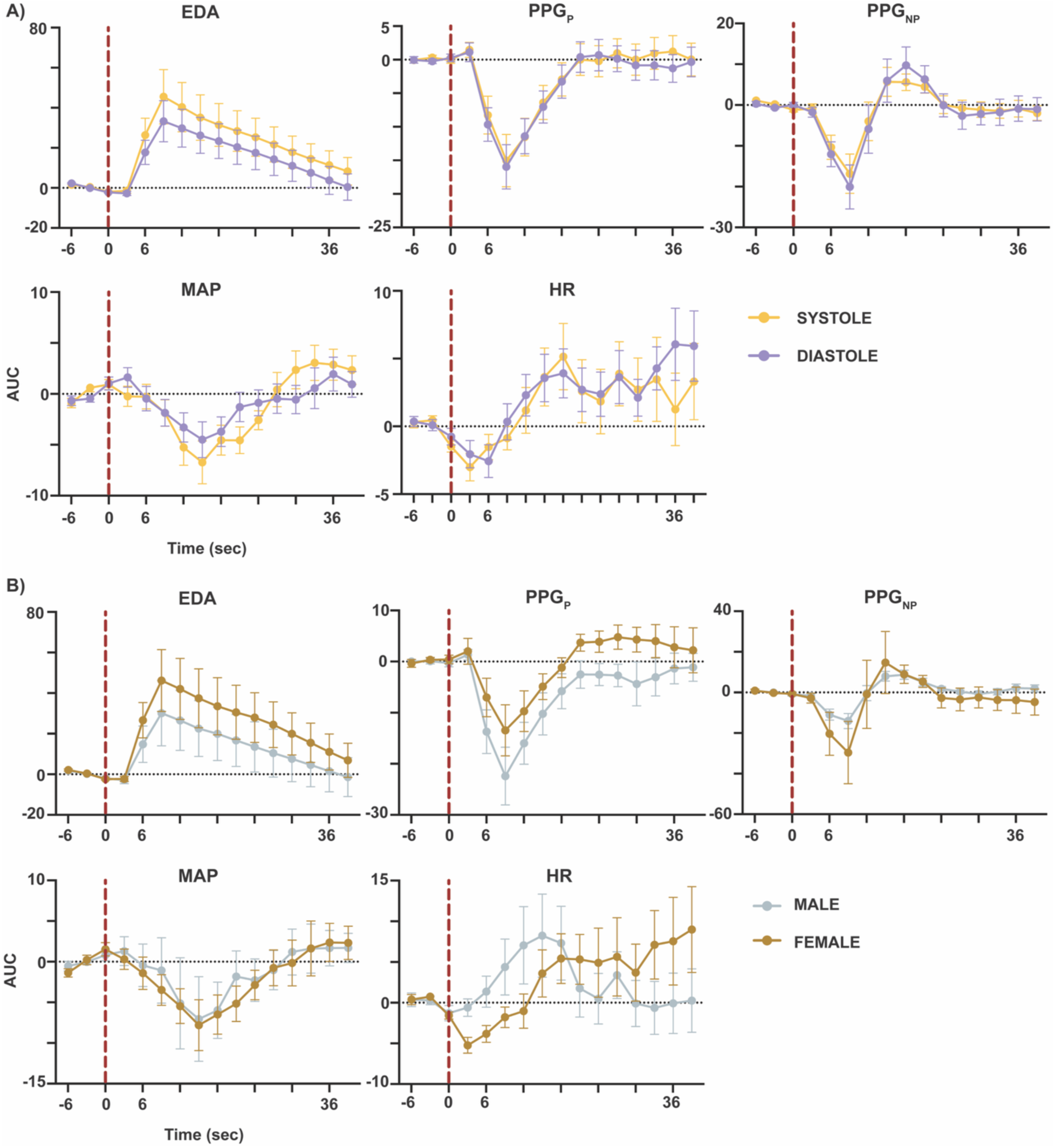
Biological Sex and Cardiac Cycle Did Not Influence Evoked Responses Following Startling Electrocutaneous Stimulation. **(A)** Evoked responses to electrocutaneous stimulation were not significantly different when stimulation was delivered either during systole (yellow traces) or diastole (purple traces). **(B)** Similarly, evoked responses to electrocutaneous stimulation were not significantly different when comparing male (gray traces) and female (beige traces) participants. Red dashed line = time of stimulation. Data are presented as mean ± SEM.

